# Mitochondrial metabolism in *Drosophila* macrophage-like cells regulates body growth via modulation of cytokine and insulin signaling

**DOI:** 10.1101/2023.02.15.528747

**Authors:** Shrivani Sriskanthadevan-Pirahas, Abdul Qadeer Tinwala, Michael J Turingan, Shahoon Khan, Savraj S Grewal

## Abstract

Macrophages play key roles in regulating and maintaining tissue and whole-body metabolism in both normal and disease states. While the cell-cell signaling pathway that underlie these functions are becoming clear, less is known about how alterations in macrophage metabolism influence their roles as regulators of systemic physiology. Here we investigate this by examining Drosophila macrophage-like cells called hemocytes. We used knockdown of TFAM, a mitochondrial genome transcription factor, to reduce mitochondrial OxPhos activity specifically in larval hemocytes. We find that this reduction in hemocyte OxPhos leads to a decrease in larval growth and body size. These effects are associated with a suppression of systemic insulin, the main endocrine stimulator of body growth. We also find that TFAM knockdown leads to decreased hemocyte JNK signaling and decreased expression of the TNF alpha homolog, Eiger in hemocytes. Furthermore, we show that genetic knockdown of hemocyte JNK signaling or Eiger expression mimics the effects of TFAM knockdown and leads to a non-autonomous suppression of body size but without altering hemocyte numbers. Our data suggest that modulation of hemocyte mitochondrial metabolism can determine their non-autonomous effects on organismal growth by altering cytokine and systemic insulin signaling. Given that mitochondrial metabolism can be controlled by nutrient availability, our findings may explain how macrophages function as nutrient-responsive regulators of tissue and whole-body physiology and homeostasis.

## Introduction

As animals develop, they need to coordinate growth across all their organs to ensure the proper attainment of functional body size. In most metazoans, this coordination relies on networks of organ-to-organ communication and endocrine signaling ^1–3^. Defects in these networks and signaling pathways can impair development leading to growth disorders and lethality.

The versatility of Drosophila genetics has led them to become a valuable model system for deciphering the tissue-to-tissue signaling networks that control body growth^2,4,5^. In Drosophila, this growth occurs during the larval stage of development and is controlled by two main endocrine systems - insulin signaling and steroid ecdysone signaling - which determine both the rate of body growth and the timing of larval maturation^6,7^. Drosophila contains eight insulins (termed Drosophila insulin-like peptides, dILPs), of which three (dILPs 2,3 and 5) are expressed and secreted from a cluster of neurosecretory cells in the brain termed insulin-producing cells (IPCs). These Dilps can circulate through hemolymph and stimulate cell, tissue and body growth by binding to a cell surface insulin receptor and activating a conserved PI3K/Akt kinase signaling pathway^8^. Ecdysone is a steroid hormone produced and secreted from the prothoracic gland (PG). Short pulses of ecdysone secretion are essential for timing the larval molts through early larval development, while a final, larger pulse of ecdysone triggers larval maturation to the pupal stage^9^. Several larval tissues can communicate with the IPC and the PG through secreted factors and cytokines to control the production and release of the dILPs and ecdysone^2^. In many cases, these tissues function as sensors of environmental factors - such as nutrition, pathogens, toxins, and oxygen - and, in turn, signal to the brain and PG to couple insulin and ecdysone production to these external changes^7^. These mechanisms of inter-organ communication allow larvae to appropriately tailor their growth and development rate to fluctuations in their environmental conditions.

Drosophila hemocytes are macrophage like-cells that can control whole-body physiology and homesotasis^10^. Like mammalian macrophages, they engulf damaged or dying cells or pathogens, most often in the context of innate immune responses^11–13^. But recent studies have emphasized their importance as regulators of organismal physiology outside of immune responses. For example, genetic depletion of hemocytes in larvae impairs growth and development and can lead to lethality^14–20^. These effects are due, in part, to reduced systemic insulin signaling and altered nutrient storage^15^. In addition, hemocyte numbers are modulated by external factors such as nutrition, oxygen levels, infection, and odorants, which may provide one way that larvae couple changes in these environmental factors to control their development and homeostasis^21,22^. For example, starvation-mediated decreases in hemocyte numbers are required for larvae to survive in poor nutrient conditions^23^. The ability of hemocytes to impact whole-body responses rely mainly on their ability to communicate with other tissues though cytokine and secreted signaling molecules.

For example, hemocytes can express and secrete upd3, a cytokine similar to mammalian interleukin-6. In larvae, hemocyte-derived upd3 can act on the fat body to suppress insulin signaling^15^ and on the PG to suppress ecdysone production and delay development^24^. In adults, hemocyte-derived upd3 mediates the impairment of systemic glucose metabolism and impaired lifespan caused by a high-fat diet^25^. Another hemocyte-secreted factor, pvf2, a fly homolog of the PDGF/VEGF growth factors, can act on the PG to suppress ecdysone production and delay larval maturation in low nutrient conditions^26^. In addition, another hemocyte-expressed pvf ligand, pvf3, can signal to the fat body to control lipid storage^27^. These findings in Drosophila parallel those from mice where tissue-resident macrophages have been shown to influence both local and whole-body systemic metabolism^28–32^.

These studies in flies and mammals emphasize the critical role that macrophages play in maintaining tissue and whole-body homeostasis beyond their phagocytic roles in immune responses to infection and tissue damage. However, little is known about macrophage metabolic responses that are important for regulating these systemic effects. Recent studies on immunity and infection in mice have shown that mitochondrial metabolic reprogramming of macrophages can determine their cytokine expression and immune responses^33^. For example, activated macrophages use mitochondrial-derived metabolites such as succinate and citrate to control the expression of interleukins and cytokines to mediate their inflammatory and immune roles^34,35^. In this paper, we have explored whether mitochondrial metabolism in Drosophila hemocytes impacts their proliferation and effects on systemic physiology and growth.

## Results

### Lowering OxPhos activity in hemocytes leads to reduced hemocyte proliferation

To explore the effects of altered mitochondrial activity on hemocyte function, we used RNAi to knock down the mitochondrial transcription factor A, TFAM. TFAM is a nuclear-encoded transcription factor that localizes to mitochondria to transcribe the mitochondrial genome, including essential components of the electron transport chain. As a result, TFAM knockdown leads to reduced mitochondrial gene expression and OxPhos activity^36,37^. We used the hemocyte-driver *hml-Gal4* to direct *UAS-TFAM RNAi* transgenes specifically in the hemocytes. We found that TFAM knockdown reduced the intensity of mitoTracker Red staining in isolated hemocytes, indicating reduced mitochondrial activity (**Fig. 1A and 1B**). We also saw that TFAM knockdown led to a significant reduction in hemocyte numbers, an effect seen with two independent RNAi lines (**Fig. 1C, 1D and Fig S1A**). In addition, TFAM knockdown prevented the increased hemocyte proliferation seen with expression of an activated form of Raf kinase, a component of the oncogenic Ras signaling pathway (**Fig. 1E**). Taken together, our results demonstrate that lowering mitochondrial bioenergetic activity through TFAM knockdown in hemocytes can suppress their proliferation even in the presence of activated Ras signaling.

**Figure 1.**
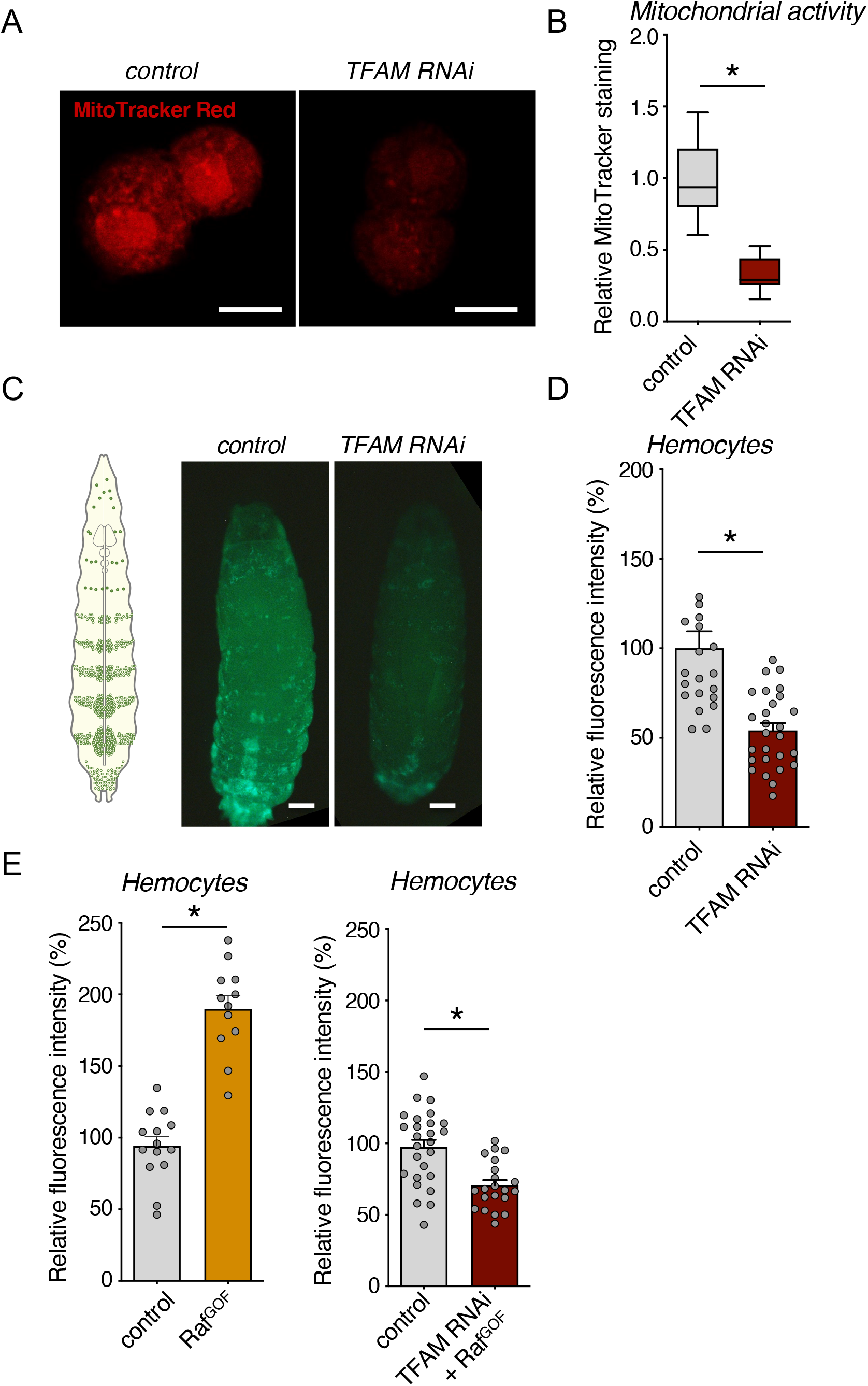
Low bioenergetic activity in hemocytes leads to reduced hemocyte proliferation. (**A**) Representative confocal micrographs of hemocyte mitochondria from control (*hml > +*) versus TFAM RNAi (*hml > UAS-TFAM-RNAi*) larvae at 96 hrs AEL stained with MitoTracker Red. The scale bars indicate 10 μm. (**B**) Quantification of MitoTracker Red staining intensity in (A). Data are presented as box plots (25%, median and 75% values) with error bars indicating the min and max values (*p < 0.05, unpaired t-test). n (# of samples) = 21 (control) and 25 (TFAM-RNAi). (**C**) Representative images of hemocytes labeled with GFP from control (*hml > +*) versus TFAM RNAi (*hml > UAS-TFAM-RNAi*) larvae at wandering stage (~144 h AEL). The scale bars indicate 100 μm. (**D**) Quantification of GFP fluorescent intensity in (C). Data are represented as mean ± SEM, with individual data points plotted as symbols (*p < 0.05, unpaired t-test). n (# of samples) = 21 (control) and 27 (TFAM-RNAi). See also Figure S1A. (**E**) Quantification of relative fluorescent intensity of GFP-labelled hemocytes in control (*hml >+*) versus Raf^GOF^ (*hml > UAS-Raf^GOF^*) and control (*hml>+*) versus Raf^GOF^ combined with TFAM-RNAi (*hml > UAS TFAM-RNAi, UAS-Raf^GOF^*). Data are represented as mean ± SEM, with individual data points plotted as symbols (*p < 0.05 and ns, not significant, unpaired t-test). n (# of samples) = 14 (control) vs 12 (Raf^GOF^) and 26 (control) vs 21 (TFAM-RNAi + Raf^GOF^).

### Hemocyte TFAM knockdown suppresses whole-body growth and development

Previous studies have shown that changes in hemocyte function can impact whole-body physiology. We, therefore, examined whether alterations in mitochondrial function bioenergetic activity might be important in these non-autonomous roles of hemocytes. We began by examining the effects on growth and development. We used RNAi to knock down TFAM in hemocytes and measured both time to pupation (as a measure of developmental rate) and pupal volume (as a measure of body size). We found that hemocyte TFAM knockdown using two independent RNAi lines led to a significant ~15-20% reduction in pupal volume (**Fig. 2A**). No effect on pupal volume was seen with transgenic flies carrying UAS-TFAM RNAi transgene alone (**Fig. S1B and S1C**). When we measured development timing, we saw a significant but minimal decrease in time to pupation (~3-6 hours) in animals with hemocyte TFAM knockdown (**Fig. 2B)**, which is unlikely to explain the substantial reduction in pupal size. Here, our results demonstrate that lowering mitochondrial bioenergetic activity in the larval hemocytes can likely suppress body size by a reduction in overall larval growth rate.

**Figure 2.**
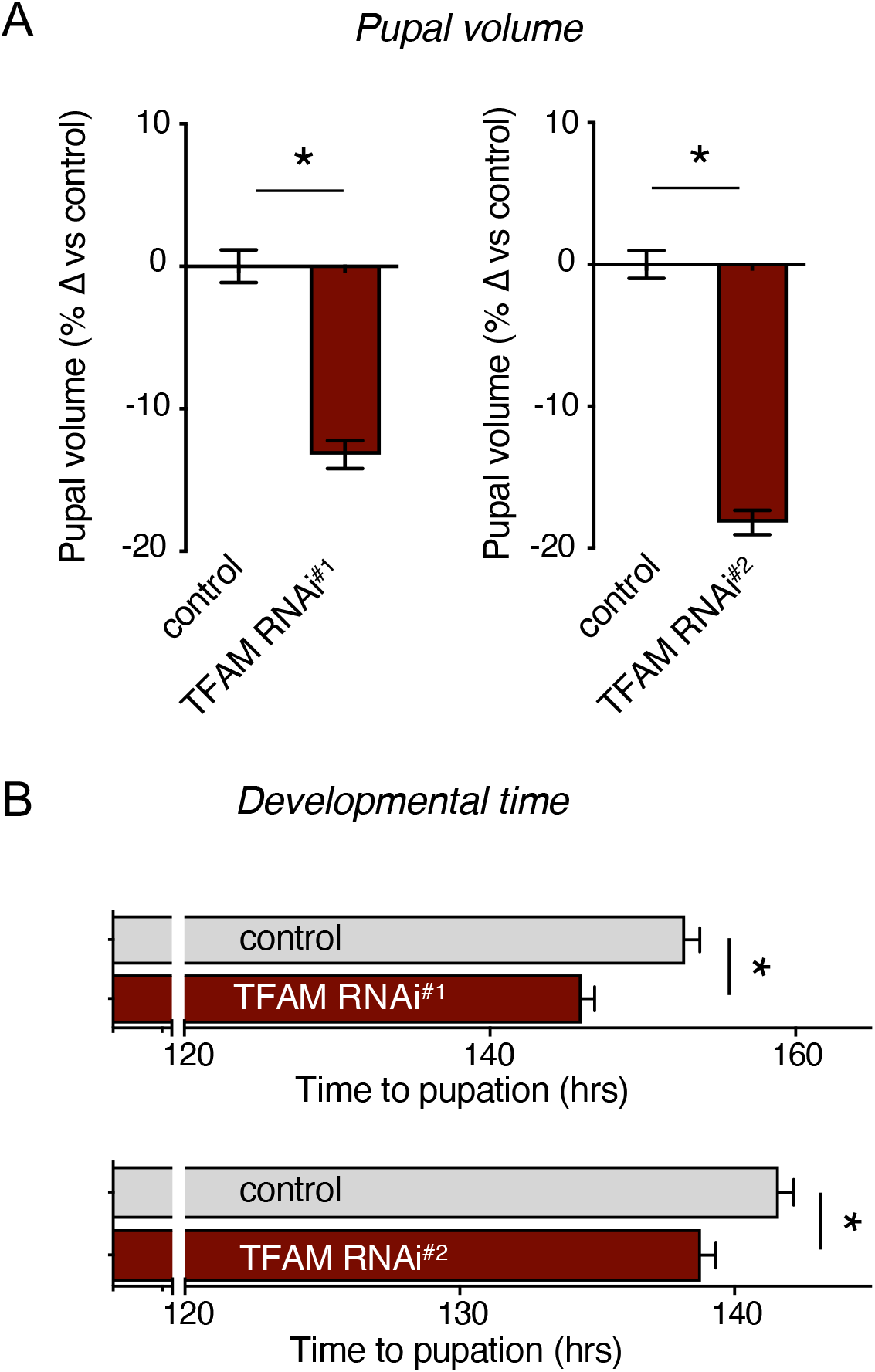
Hemocyte TFAM knock down suppresses systemic growth and development. (**A**) Relative change in pupal volume was calculated based on the average value of control (hml>+) animals. Data are presented as mean +/− SEM (*p < 0.05, Mann-Whitney U test) for controls and two different TFAM RNAi lines (*hml > TFAM-RNAi*). n (# of pupae) = 183 (control) vs 195 (TFAM RNAi#^1^) and 199 (control) vs 179 (TFAM RNAi#^2^). See also Figure S1B. (**B**) Time to pupation was measured in control (*hml > +*) larvae versus larvae expressing one of two different TFAM RNAi transgenes (*hml > UAS-TFAM RNAi*). Data are presented as mean time to pupation +/− SEM (*p < 0.05, Mann-Whitney U test). n (# of pupae) = 171 (control) vs 135 (TFAM RNAi#^1^) and 179 (control) vs 107 (TFAM RNAi#^2^). See also Figure S1C.

### Hemocyte TFAM knockdown suppresses systemic insulin signaling

We next explored how hemocyte TFAM knockdown might suppress overall body growth. One of the central regulators of growth in larvae is the endocrine insulin pathway. Flies have seven Drosophila insulin-like peptides (dILPs). These bind to a single inulin receptor and activate a conserved PI3 Kinase/Akt kinase signaling pathway that can stimulate growth in all larval tissues. We found that hemocyte knockdown of TFAM led to reduced levels of whole-body phosphorylated Akt (**Fig. 3A and 3B**), consistent with reduced systemic insulin signaling. One primary way insulin signaling is controlled is through the production and release of three dILPS (2, 3 and 5) from the insulin-producing cells (IPCs) in the brain^8^. We found that hemocyte TFAM knockdown did not affect the whole-body mRNA levels of any of the seven dILPs (**Fig. S2A**). However, when we used dILP2 antibody staining to examine the IPCs, we saw an accumulation of dILP protein (**Fig. 3C and 3D**), an effect characteristic of dILP2 retention due to reduced dILP2 secretion^38^. Taken together, our results suggest that one way that hemocyte-specific knockdown of TFAM leads to reduced body growth is by lowering brain-derived dILP secretion leading to suppressed systemic insulin signaling. One well-described target of the insulin/PI3K/Akt pathway is the transcription factor FOXO, whose nuclear localization and transcriptional activity are usually inhibited by Akt. Interestingly, we found that hemocyte-specific TFAM RNAi didn’t affect the whole-body expression levels of known FOXO target genes, 4E-BP, dILP6, and InR^39,40^ (**Fig. S2B**). Furthermore, when we stained fat body tissue for FOXO, we observed a substantial decrease in total FOXO protein levels rather than any change in nuclear vs cytoplasmic localization **(Fig. 4A and 4B)**. Levels of *foxo* mRNA were unchanged. Previous work showed that *foxo* mull mutants had reduced final adult tissue and body size^41^. We found that *foxo* mutants had reduced pupal volume, and this size reduction was not further exacerbated by hemocyte TFAM knockdown **(Fig 4C)**. These results show that lowering hemocyte OxPhos activity by TFAM knockdown can decrease systemic insulin signaling and reduce FOXO protein levels. Both effects may explain the decrease in whole-body growth.

**Figure 3.**
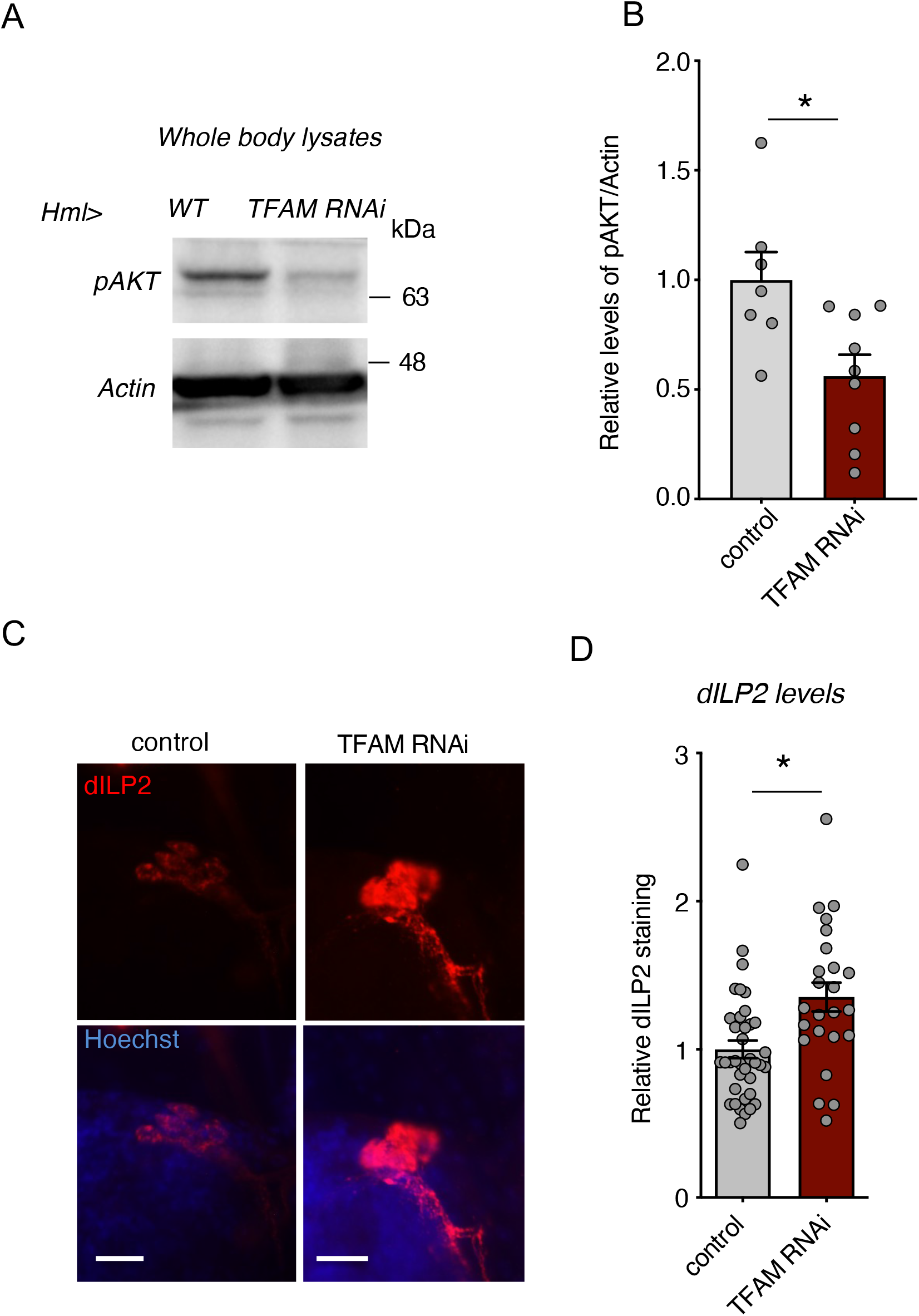
Hemocyte TFAM knock suppresses systemic insulin signaling by inhibiting dILP2 secretion from brain IPCs. (**A**) Western blots of whole-body samples from control (*hml > +*) versus TFAM RNAi (*hml > UAS-TFAM-RNAi*) larvae at 96hrs AEL analyzed using Phospho-Akt and actin antibodies. (**B**) Quantification of western blots from (A). Data are relative levels of phospho-Akt band intensity corrected for actin band intensity. Data are presented as box plots (25%, median and 75% values) with error bars indicating the min and max values (*p < 0.05 unpaired t-test, n = 7 (control) and 9 (TFAM RNAi) groups per condition with 20 larvae in each group). (**C**) Representative images for brain IPCs stained with dILP2 in control (*hml > +*) versus TFAM RNAi (*hml > UAS-TFAM-RNAi*) larvae at 96 hrs AEL larvae. The scale bars indicate 20 μm. (**D**) Quantification of relative dILP2 fluorescent intensity in (C). Data are represented as mean ± SEM, with individual data points plotted as symbols (*p < 0.05, unpaired t-test). n (# of samples) = 37 (control) and 24 (TFAM-RNAi).

**Figure 4.**
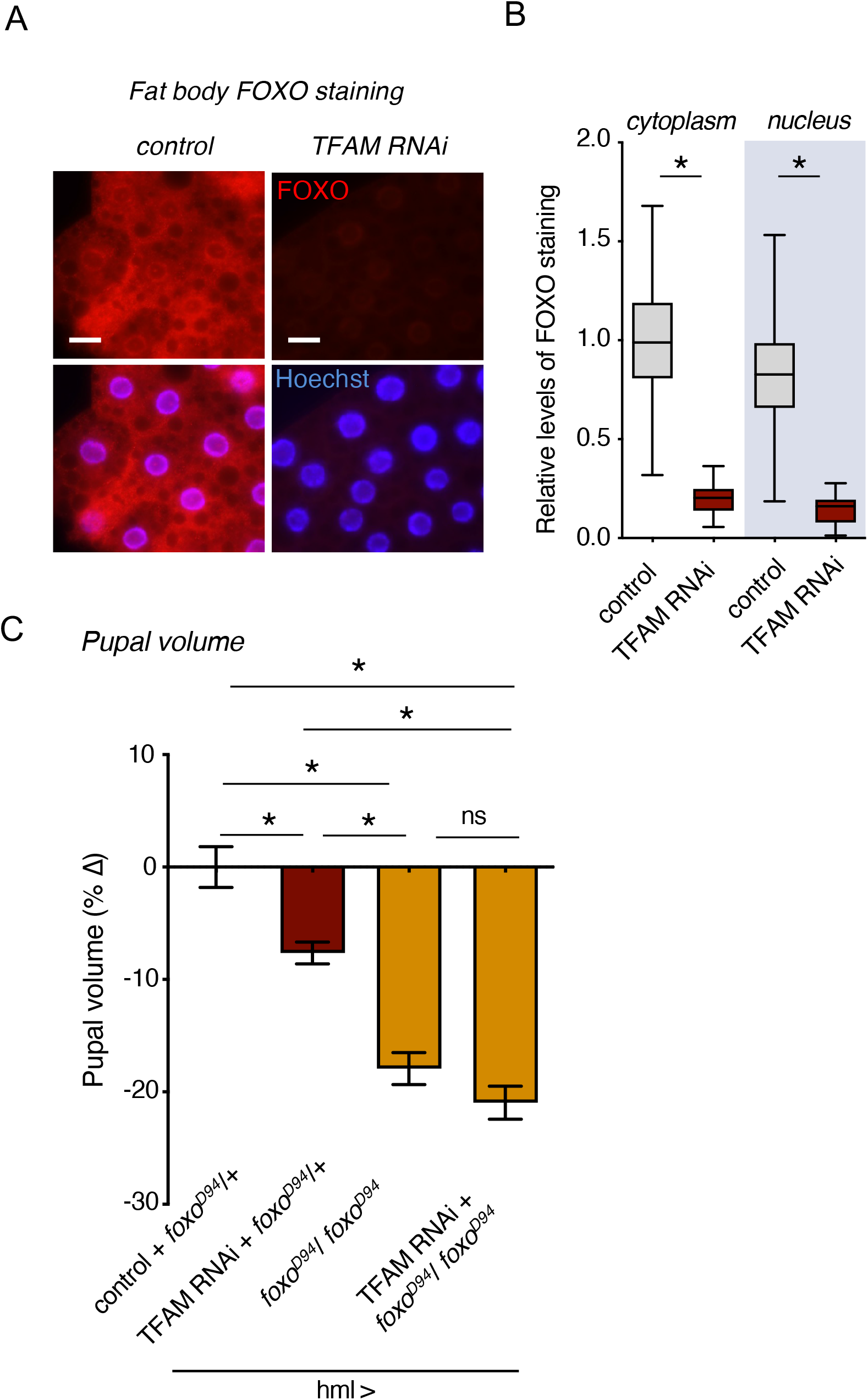
Hemocyte TFAM knock down leads to suppression of fat body FOXO levels. (**A**) Representative images for fat body stained with anti-FOXO antibodies from control (*hml > +*) versus TFAM RNAi (*hml > UAS-TFAM-RNAi*) larvae at 96 hrs AEL larvae. The scale bars represent 20 μm. (**B**) Quantification of relative FOXO fluorescent intensity in the fat body cytoplasm and nucleus (C). Data are presented as box plots (25%, median and 75% values) with error bars indicating the min and max values (*p < 0.05, unpaired t-test). n (# of fat cells) = 128 (control) and 124 (TFAM-RNAi). (**C**) Relative change in pupal volume was calculated based on the average value of control (hml>+) animals. Data are presented as mean +/− SEM (*p < 0.05 and ns, not significant, Mann-Whitney U test) for control (*hml > +, foxo^Δ94^/+*), TFAM RNAi (*hml > UAS-TFAM-RNAi, foxo^Δ94^/+*), *foxo^Δ94^* (*hml> +, foxo^Δ94^/foxo^Δ94^*), and TFAM-RNAi + *foxo^Δ94^* (*hml > UAS-TFAM RNAi, foxo^Δ94^/foxo^Δ94^*). n (# of pupae) = 73 (control), 224 (TFAM RNAi), 107 (*foxo^Δ94^*), and 97 (TFAM RNAi + *foxo^Δ94^*).

### TFAM knockdown inhibits hemocyte JNK signaling pathway, and genetic suppression of hemocyte JNK signaling suppresses body growth

We next wanted to investigate the downstream effects of TFAM knockdown in hemocytes. Alterations in mitochondrial function have been shown to modulate JNK signaling pathway activity in Drosophila^42^. We there examined phosphorylation levels of the JNK. We found hemocytes with TFAM knockdown have significantly lower pJNK staining (**Fig. 5A and 5B**), suggesting that lowering mitochondrial OxPhos suppresses the JNK signaling pathway. To explore whether this decrease in JNK signaling might explain the effects of hemocyte TFAM knockdown on both hemocyte proliferation and body growth, we genetically inhibited JNK pathway activity in hemocytes by expression of either a dominant negative version of the JNK kinase, *Basket* (*BskDN*), or by RNAi-mediated knockdown of *Kayak* (*Kay RNAi*), a transcriptional target and effector of JNK signaling in flies. In both cases, we saw that inhibition of JNK signaling in hemocytes had no effects on hemocyte numbers **(Fig. 5C and 5E),** suggesting that suppression of JNK signaling does not explain the reduced hemocyte proliferation following TFAM knockdown. However, we saw that hemocyte-specific expression of either *BskDN* or hemocyte *Kay RNAi* led to a reduction in pupal volume **(Fig. 5D and 5F**) and that *Kay RNAi* had little effect on time to pupation **(Fig S3A).** In addition, we saw that the reduction in body size following hemocyte-specific knockdown of TFAM and Kay was comparable to the effects of either knockdown alone, suggesting both factors function similarly **(Fig 5F)**. Together, our results suggest that one way that the reduction of OxPhos activity by TFAM knockdown suppresses whole-body growth through reduced activity of the JNK-signaling pathway and that these effects are independent of any changes in hemocyte number.

**Figure 5.**
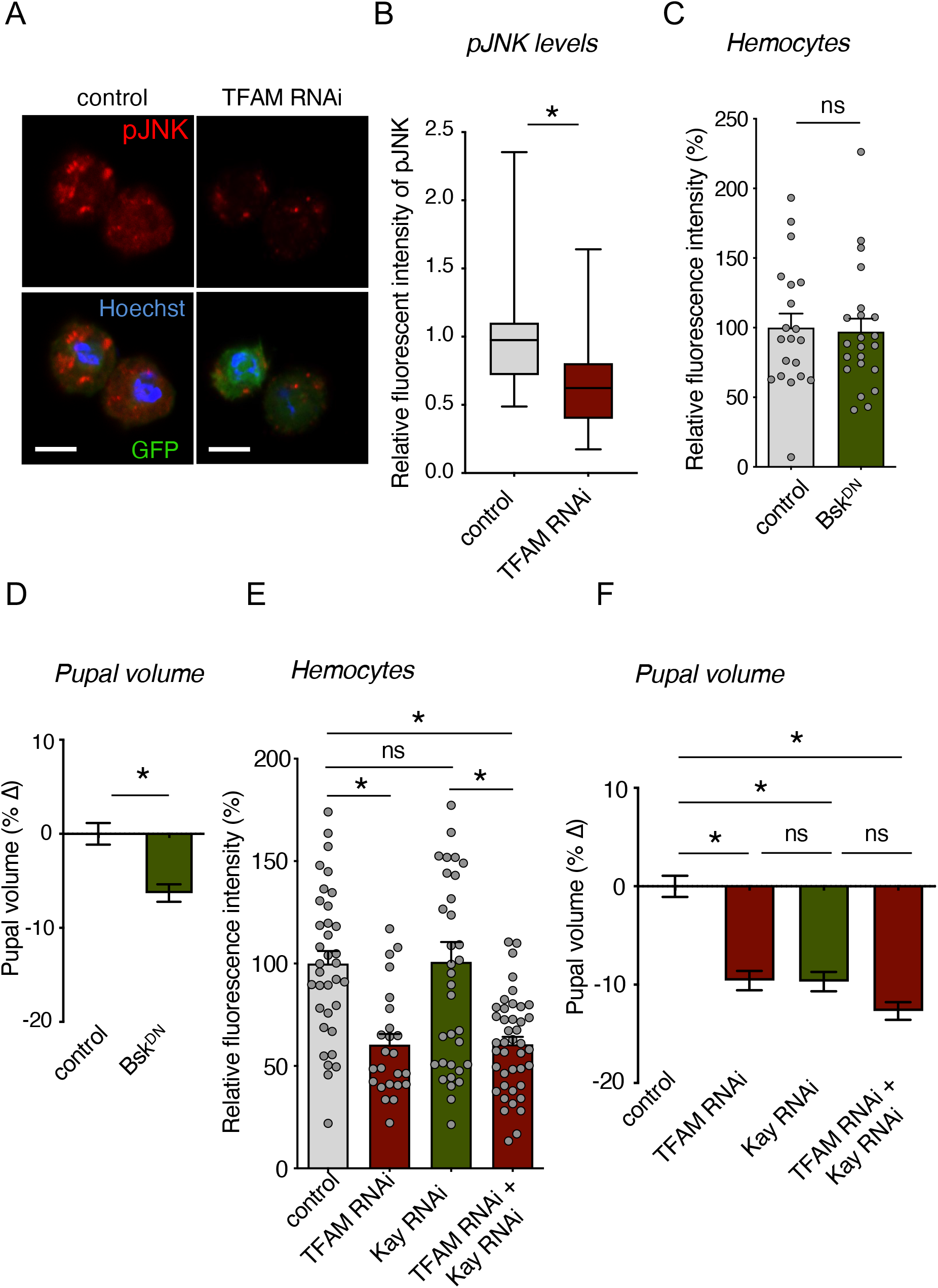
Hemocyte specific knock down of JNK signaling suppresses systemic growth. (**A**) Representative images for hemocytes stained with phospho-JNK (pJNK) in control (*hml > +*) versus TFAM RNAi (*hml > UAS-TFAM-RNAi*) larvae at 96 hrs AEL larvae. The scale bars represent 5 μm. (**B**) Quantification of relative pJNK fluorescent intensity in hemocytes (A). Data are presented as box plots (25%, median and 75% values) with error bars indicating the min and max values (*p < 0.05, unpaired t-test). n (# of hemocytes) = 36 (control) and 36 (TFAM-RNAi). (**C**) Quantification of relative fluorescent intensity of GFP-labelled hemocytes in control (*hml > +*) versus Bsk^DN^ (*hml > Bsk^DN^*). Data are represented as mean ± SEM, with individual data points plotted as symbols (*p < 0.05 and ns, not significant, unpaired t-test). n (# of samples) = 20 (control) and 22 (Bsk^DN^). (**D**) Relative change in pupal volume was calculated based on the average value of control (*hml > +*) animals. Data are presented as mean +/− SEM (*p < 0.05 and ns, not significant, Mann-Whitney U test) for control (*hml > +*) vs Bsk^DN^ (*hml > BskD^N^*) animals. n (# of pupae) = 206 (control), 206 (Bsk^DN^). (**E**) Quantification of relative fluorescent intensity of GFP-labelled hemocytes in control (*hml > +*), TFAM RNAi (*hml > UAS-TFAM RNAi*), Kay RNAi (*hml > UAS-Kay RNAi*) and TFMA RNAi + Kay RNAi (*hml > UAS-TFAM RNAi + UAS-Kay RNAi*) larvae. Data are represented as mean ± SEM, with individual data points plotted as symbols (*p < 0.05 and ns, not significant, unpaired t-test). n (# of samples) = 35 (control), 35 (*TFAM RNAi*), 24 (*Kay RNAi*), and 42 (*TFAM RNAi + Kay RNAi*). (**F**) Relative change in pupal volume was calculated based on the average value of control (hml>+) animals. Pupal volume data analysis in control (*hml > +*), TFAM RNAi (*hml > UAS-TFAM RNAi*), Kay RNAi (*hml > UAS-Kay RNAi*) and TFMA RNAi + Kay RNAi (*hml > UAS-TFAM RNAi + UAS-Kay RNAi*) larvae. Data are represented as mean ± SEM, with individual data points plotted as symbols (*p < 0.05 and ns, not significant, unpaired t-test). n (# of samples) = 180 (control), 202 (*TFAM RNAi*), 213 (*Kay RNAi*), and 200 (*TFAM RNAi + Kay RNAi*).

### Hemocyte Eiger expression is controlled by mitochondrial OxPhos and can regulate body size

Hemocytes express many different secreted factors that can impact whole-body physiology. Interestingly, we observed a significant reduction in hemocyte mRNA levels of one such factor – the TNF alpha homolog, Eiger – following TFAM knockdown in hemocytes (**Fig. 6A**). We also saw that RNAi-mediated Eiger knockdown in hemocytes led to smaller body size (**Fig. 6B**) without any effect on hemocyte number (**Fig. 6C**). Flies carrying just the UAS-Eiger RNAi transgene alone showed no significant change in body size (**Fig. S3B)** These results suggest that one way that lowered hemocyte mitochondrial OxPhos activity suppresses body growth is through altered cytokine signaling.

**Figure 6.**
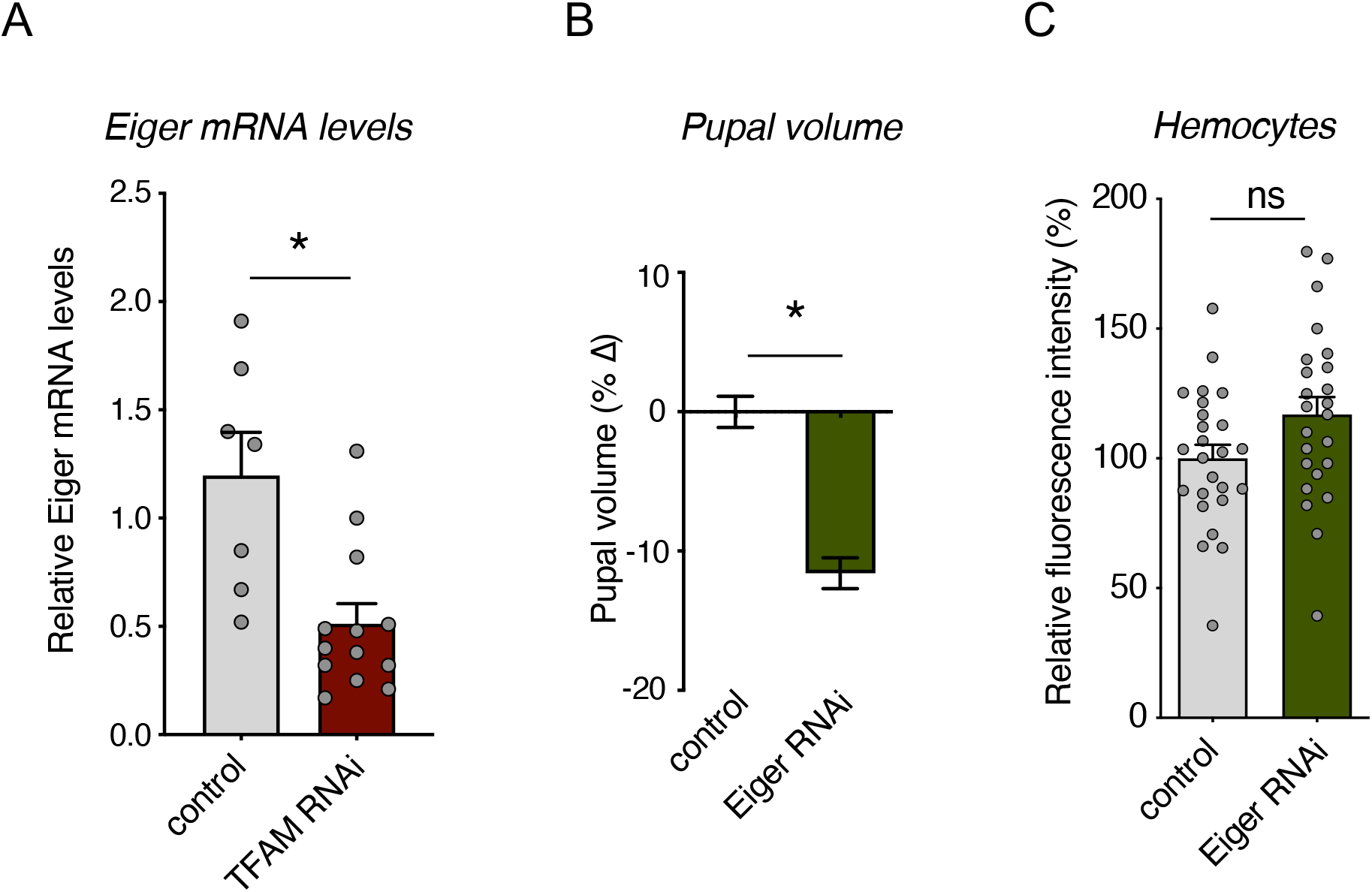
Hemocyte specific cytokine knock down suppresses body growth. (**A**) Hemocyte specific Eiger mRNA levels measured by qRT-PCR in control (*hml > +*) versus TFAM RNAi (*hml > UAS-TFAM-RNAi*) larvae at 120 hrs AEL. Data are represented as mean ± SEM, with individual data points plotted as symbols (*p < 0.05 and ns, not significant, unpaired t-test). n (# of samples) = 7 (control) and 13 (TFAM RNAi). (**B**) Relative change in pupal volume of control (*hml > +*) versus Eiger RNAi (*hml > UAS-Eiger RNAi*) larvae. Data are presented as mean +/− SEM (*p < 0.05 and ns, not significant, Mann-Whitney U test). n (# of pupae) = 187 (control), 165 (Eiger RNAi). (**C**) Quantification of relative fluorescent intensity of GFP-labelled hemocytes in control (*hml > +*) versus Eiger RNAi (*hml > UAS-Eiger RNAi*) larvae. Data are represented as mean ± SEM, with individual data points plotted as symbols (*p < 0.05 and ns, not significant, unpaired t-test). n (# of samples) = 25 (control) and 24 (Eiger RNAi).

## Discussion

Our main finding is that suppressing hemocyte mitochondrial OxPhos activity can exert both autonomous and non-autonomous effects on growth **(Fig 7)**. The autonomous effects involve a block in hemocyte proliferation even when proliferation is stimulated by activation of the oncogenic Ras pathway. This result is consistent with similar studies in mouse models of lung cancer showing that mitochondrial metabolism is essential for Ras-mediated tumors^36^. The non-autonomous effect of reduction in hemocyte OxPhos was suppression in overall body growth. The final pupal volume was reduced, but developmental timing was only modestly accelerated. Therefore, we likely saw decreased body size resulting from reduced overall growth rather than an acceleration of the growth period. Consistent with this, we saw that the decrease in body growth caused by hemocyte TFAM knockdown was accompanied by reduced whole body Akt phosphorylation and reduced dILP2 release from the brain, pointing to a reduction in systemic insulin signaling, the main regulator of body growth. Interestingly, we saw a reduction in FOXO protein levels, even though reduced insulin typically leads to FOXO nuclear accumulation. However, regulation of FOXO protein levels has been observed previously in Drosophila^43,44^. Interestingly, *foxo* null mutants have reduced body size, a phenotype that we also see here and this body size phenotype we observed was not further exacerbated by hemocyte TFAM knockdown. Therefore, hemocyte mitochondrial metabolism may control body growth by controlling both brain-derived insulin signaling and regulation of FOXO levels.

**Figure 7.**
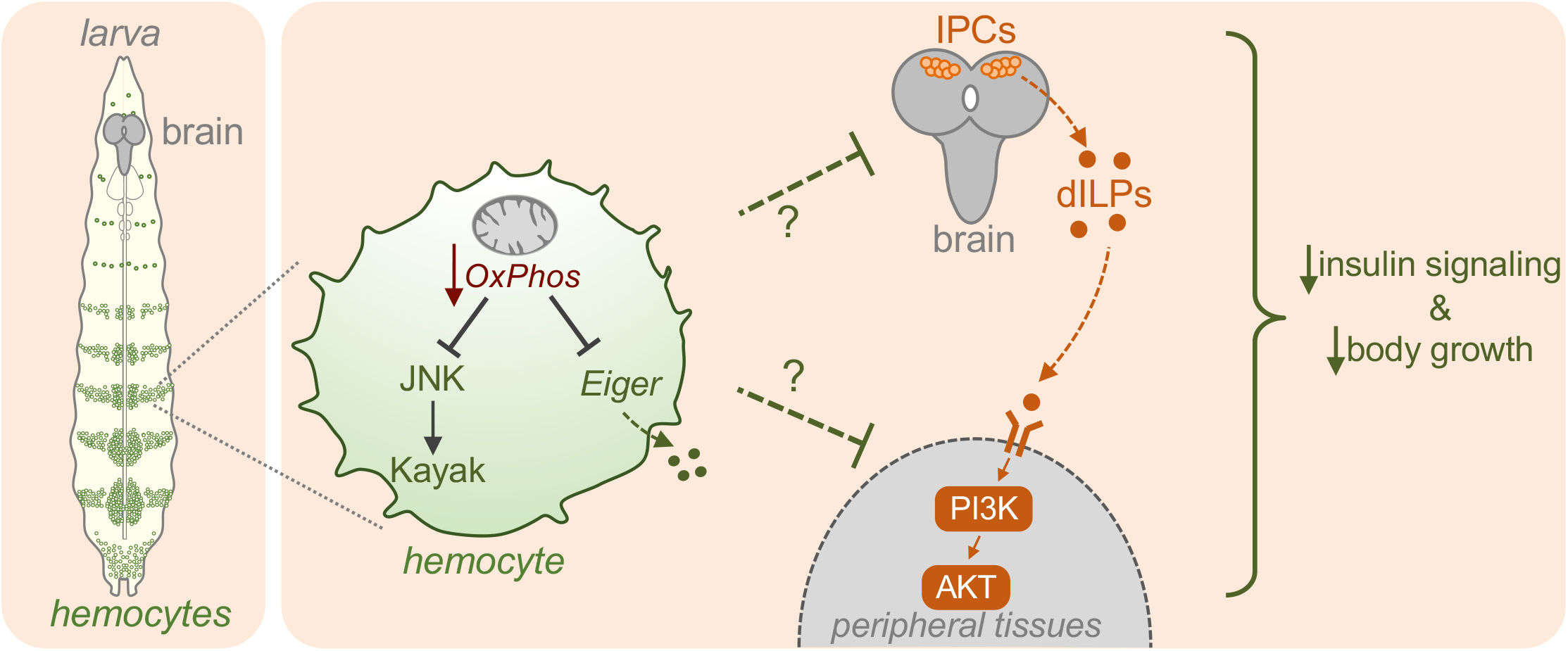
Low bioenergetic mitochondrial activity in hemocyte leads to suppression of systemic insulin signaling. When hemocyte mitochondrial OxPhos activity is low (for example following TFAM knockdown), expression of Eiger and the activity of the JNK pathway are reduced. Under these conditions, dILP2 secretion from the brain IPC cells and systemic insulin signaling are reduced leading to reduced animal growth and development.

One potential effector of reduced OxPhos activity in hemocytes is the reduction of JNK signaling. We saw that TFAM knockdown leads to reduced phosphorylation of JNK and that genetic suppression of JNK signaling in hemocytes could mimic the effects of TFAM knockdown reduced body size. Interestingly, the effects of JNK pathway suppression and TFAM knockdown on body size were not additive, suggesting they function in the same pathway. Our work also identified the hemocyte-expressed secreted factor, Eiger, as a potential link between changes in hemocyte OxPhos activity to control of body growth. We also saw that TFAM knockdown decreased Eiger mRNA expression in hemocytes and that hemocyte knockdown of Eiger mimicked the effects of TFAM knockdown and led to a reduction in body size. These results suggest that reduced hemocyte OxPhos activity may decrease Eiger expression to mediate non-autonomous effects on body size. Interestingly, fat derived Eiger can inhibit dILP secretion from IPCs ^45^. Hence it may appear paradoxical that blocking hemocyte production of Eiger would lead to reduced body size. However, it is possible that the effects on body size we see with Eiger manipulation either occur independently of changes in systemic insulin signaling, or that the effects of Eiger on systemic insulin signaling may depend on the cell type it is expressed from (fat vs hemocytes).

An interesting finding from our work was that the body size suppression caused by hemocyte suppression of JNK signaling or Eiger knockdown was independent of any change in hemocyte number. This suggests that the non-autonomous effects of hemocyte OxPhos reduction on body growth may not be because of changes in hemocyte number. Similarly, a recent report showed that the effects of hemocytes on developmental timing mediated by pvf2 signaling in response to low nutrients were also independent of hemocyte cell number^26^. In contrast, another study reports that a reduction in hemocyte numbers is needed for their effects on nutrient storage and survival in poor nutrients^23^. Thus, both the metabolic status and numbers of hemocytes are important for determining their impact on organismal physiology.

A question prompted by our work is how does a reduction in hemocyte mitochondrial OxPhos activity lead to suppression of JNK signaling and Eiger expression? One possibility is that these effects are caused by alterations in reactive oxygen species (ROS) levels. The mitochondrial electron transport chain is a major source of ROS production in cells, and JNK is stimulated by ROS levels^46,47^. Thus, TFAM knockdown, by lowering OxPhos activity, may limit ROS production and thereby reduce JNK activity. Additionally, a reduction in OxPhos may reprogram mitochondrial metabolism leading to alterations in TCA cycle intermediates. Changes in the levels of these metabolites, such as succinate and citrate, have previously been shown to couple mitochondrial metabolism in activated mammalian macrophages to the expression of cytokines by altering the activity of chromatin modifiers^33,48^. Hence, a similar mechanism may regulate Eiger expression in hemocytes.

Macrophages play multiple essential roles in regulating tissue and whole-body metabolic homestasis^28–32^. These general regulatory roles of macrophages are seen in both invertebrates and vertebrates and are often influenced by changes in nutrients^27,29^. Given that regulation of mitochondrial metabolism is a downstream target of many conserved nutrient-responsive signaling pathays^49^, our findings suggest that changes in mitochondrial metabolism may link the nutrient-sensing properties of macrophages to their role as regulators of metabolic homeostasis.

## Methods

### Drosophila food and genetics

Flies were raised on a medium containing 150 g agar, 1600 g cornmeal, 770 g Torula yeast, 675 g sucrose, 2340 g D-glucose, 240 ml acid mixture (propionic acid/phosphoric acid) per 34 L water and maintained at 25 °C. For all GAL4/UAS experiments, homozygous GAL4 lines were crossed to the relevant UAS line(s) and the larval progeny were analyzed. Control animals were obtained by crossing the appropriate homozygous GAL4 line to flies of the same genetic background as the relevant experimental UAS transgene line.

### Drosophila Strains

The following strains were used: *w^1118^, hml-GAL4, UAS-GFP, UAS-Eiger RNAi* (VDRC 108814), GD control line (60000 TK), KK control line (VDRC 60100 TK), UAS-Kayak-RNAi (VDRC 6212 GD), *UAS-TFAM RNAi #2 (4217R-2), UAS-TFAM RNAi #3 (4217R-3 - Fly Stocks of National Institute of Genetics - NIG-FLY), foxo^Δ94^*(gift from Linda Partridge)^41^.

### Measurement of *Drosophila* developmental time

For measuring development timing to the pupal stage, newly hatched larvae were collected at 24 hrs AEL and placed in food vials (50 larvae per vial). The number of newly formed pupae was counted twice a day until all larvae had pupated.

### Pupal imaging and pupal volume measurement

Pupae were imaged using a Zeiss Discovery V8 Stereomicroscope with Axiovision imaging software. Pupal length and width were measured, and pupal volume was calculated using the formula, volume=4/3π(L/2) (l/2)2.

### Quantification of hemocyte number

To assay for hemocyte numbers, we used an *hml-GAL4, UAS-GFP*, to GFP-label hemocytes. Larvae were collected during the L3 wandering larval stage using forceps and cleaned by being placed in a small petri dish containing 5 mL of phosphate buffered saline (PBS) for 30 seconds. The clean larvae were then transferred to another small petri dish and fluorescence-imaged using ZEISS SteREO Discovery V8 microscope and ZEN imaging software (blue edition) at 8.0x magnification. Next, the NIH ImageJ software was used to quantify the fluorescence intensity in a defined region in posterior segments of each larvae where hemocytes are clustered. This value was then corrected for background autofluorescence by subtracting the average fluorescence intensity measured from unlabeled *w^1118^* L3 wandering larvae.

### MitoTracker Red staining

Hemocytes from 96hrs AEL larvae were collected and stained with MitoTracker Deep Red FM (1: 1000 dilution of 1 mM, Molecular probes M22426) for 40 mins and fixed at room temperature using 8% PFA for 30 mins. After washing three times, mounted using Vecta Shield mounting medium. The mitochondrial images were acquired through Zeiss confocal microscope LSM 880.

### Preparation of larval protein extracts

*Drosophila* larvae (96 hrs. AEL) were lysed with homogenization and sonication in a buffer containing 20 mM Tris-HCl (pH 8.0), 137 mM NaCl, 1 mM EDTA, 25% glycerol, 1% NP-40 and with the following inhibitors: 50 mM NaF, 1 mM PMSF, 1 mM DTT, 5 mM sodium ortho vanadate (Na3VO4) and Protease Inhibitor cocktail (Roche Cat. No. 04693124001) and Phosphatase inhibitor (Roche Cat. No. 04906845001), according to the manufacturer instructions.

### Western blots, immunostaining and antibodies

Protein concentrations were measured using the Bio-Rad Dc Protein Assay kit II (Bio-Rad 5000112). Protein lysates (100 μg) were resolved by SDS-PAGE and electro transferred to a nitrocellulose membrane, subjected to western blot analysis with specific antibodies, and visualized by chemiluminescence (enhanced ECL solution (Perkin Elmer)). The primary antibodies used in this study were: anti-phospho-AKT-Ser505 (1:1000, Cell Signaling Technology #4054), anti-actin (1:1000, Santa Cruz Biotechnology, # sc-8432), anti-dILP2^50^ (1:500), and anti-phospho-JNK (1:500, Cell Signaling Technolog #4668S). Goat and donkey secondary antibodies were purchased from Santa Cruz Biotechnology (sc-2030, 2005, 2020). The rabbit anti-FOXO antibody was used at 1:500 dilution for fat body immunostaining (a gift from Marc Tatar).

### Larval brain staining

Larval brains were dissected and fixed for 30 min in 4% formaldehyde in PBS, washed three times in PBS with 0.1% Triton X-100 (PBT). Tissues were then pre-blocked in PBT + 5% BSA + 2% fetal bovine serum for 2 hrs and then incubated overnight at 4°C with the primary antibody (1:1000 dilution of anti-dILP2) in 5% BSA+ PBT, and then washed three times in PBT+0.5% BSA. A cocktail of secondary antibodies was then added to the block (final concentration 1:400) and the tissues were incubated overnight in secondary at 4°C. The samples were then washed 3× for 15 min each time, with PBT+0.5% BSA and mounted on slides. Brains were imaged using Zeiss Stereo Discovery V8 microscope using Axiovision software.

### Quantitative RT-PCR measurements

Total RNA was extracted from hemocytes collected from 20 larvae at 120h AEL or from total larval lysates at 96h AEL using TRIzol according to manufacturer’s instructions (Invitrogen; 15596-018). RNA samples isolated from the same number of larvae (control vs experimental) were DNase treated (Ambion; 2238G) and reverse transcribed using Superscript II (Invitrogen; 100004925). The generated cDNA was used as a template to perform qRT–PCRs (ABI 7500 real time PCR system using SyBr Green PCR mix) using gene-specific primers. PCR data were normalized to RpL32. All primer sequences are listed in the supplementary table S1.

### Statistical analysis

All qRT-PCR data and quantification of immunostaining data were analyzed by Students t-test, two-way ANOVA followed by post-hoc students t-test, or Mann-Whitney U test where appropriate. All statistical analysis and data plots were performed using Prism statistical software. Differences were considered significant when p values were less than 0.05.

## Acknowledgements

We thank Linda Partridge and Kim Rewitz for the gift of fly stocks and antibodies. Stocks obtained from the VDRC, the NIG-Fly Stock Centre, Kyoto, Japan, and the Bloomington Drosophila Stock Center (NIH P40OD018537) were used in this study. This work was supported by CIHR Project Grants (PJT-173517, PJT-152892) and Cancer Research Society grants to S.S.G. M.J.T. was supported by an NSERC summer studentship.

## Author contributions

SS-P and SSG conceived and managed the project. SS-P, AQT, MJT and SK performed experiments and analyzed data. SS-P and SG drafted and revised the manuscript with feedback from all authors.

## Data availability statement

All data needed to evaluate the conclusions in the paper are present in the paper and/or the Supplementary Materials. The datasets used and/or analysed during the current study are available from the corresponding author.

---

**Figure S1:**
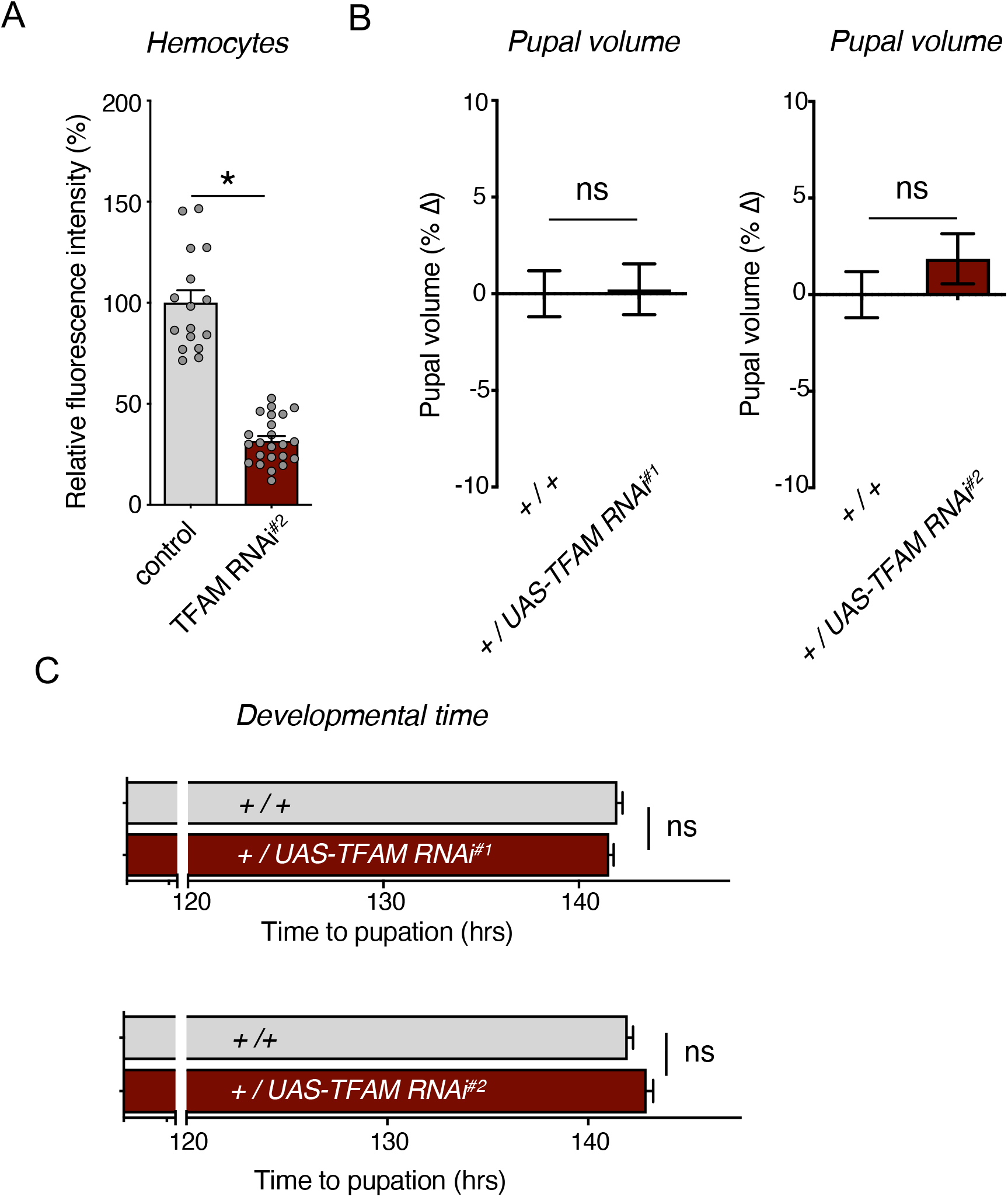
TFAM knock down in hemocytes suppresses hemocyte proliferation and systemic growth (related to Fig. 1 and 2) (A) Quantification of relative fluorescent intensity of GFP-labelled hemocytes from control (*hml>+*) versus TFAM RNAi#^2^ (*hml>UAS-TFAM-RNAi*) larvae at wandering stage. Data are represented as mean ± SEM, with individual data points plotted as symbols (*p < 0.05, unpaired t-test). n (# of samples) = 16 (control) and 23 (TFAM-RNAi^#2^). (B) Relative change in pupal volume was calculated based on the average value of control (*+/+*) animals. Data are presented as mean +/− SEM (*p < 0.05, Mann-Whitney U test) for controls and two different TFAM RNAi lines (*+/UAS-TFAM-RNAi*). n (# of pupae) = 147 (control) vs 148 (TFAM RNAi#^1^) and 147 (control) vs 146 (TFAM RNAi#^2^). (C) Time to pupation was measured in control (+/+) larvae versus larvae expressing one of two different TFAM RNAi transgenes (*+/UAS-TFAM RNAi*). Data are presented as mean time to pupation +/− SEM (*p < 0.05, Mann-Whitney U test). n (# of pupae) = 590 (control) vs 675 (TFAM RNAi#^1^) and 590 (control) vs 592 (TFAM RNAi#^2^).

**Figure S2:**
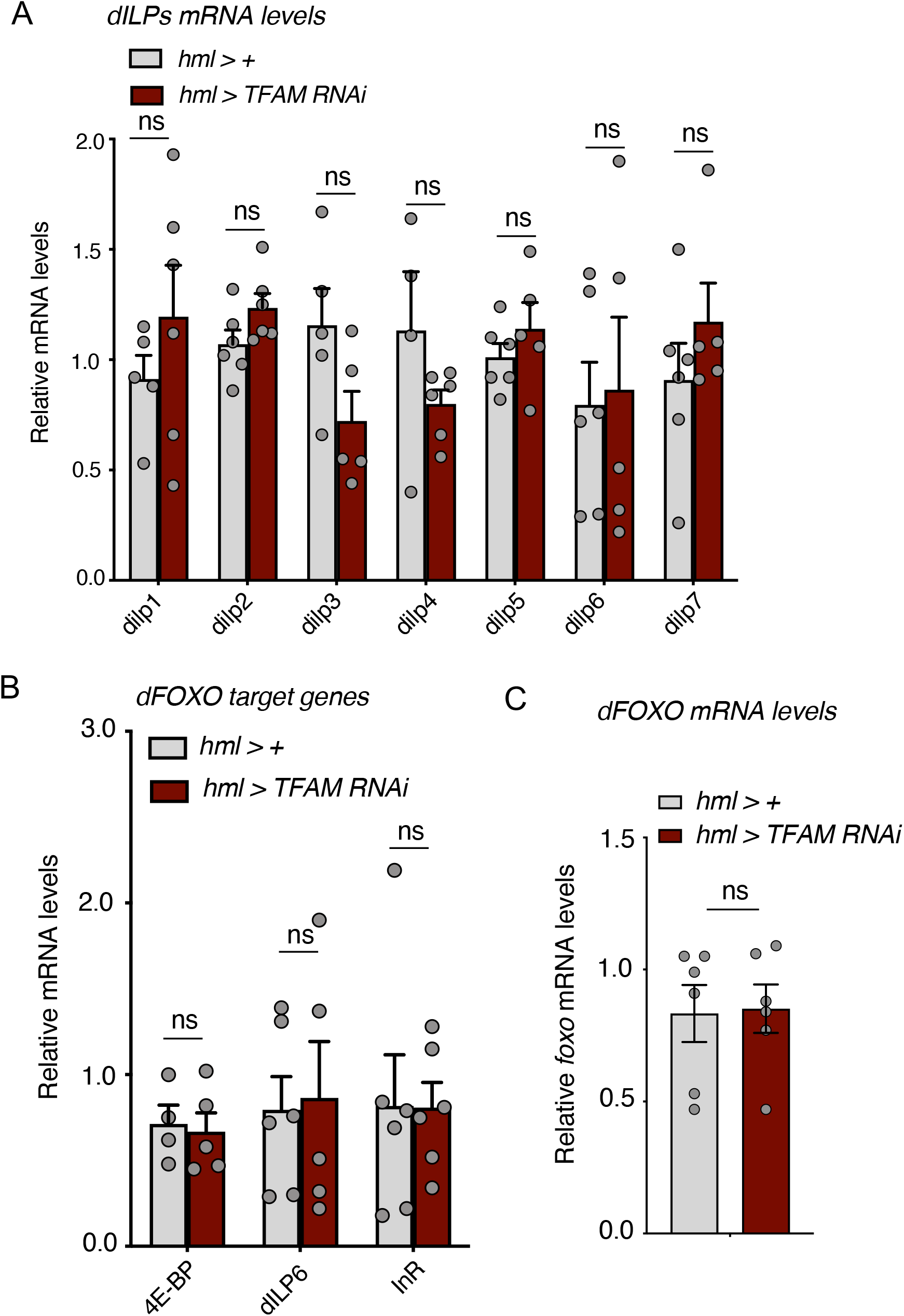
Hemocyte TFAM knock down shows no change in whole larvae mRNA levels of *dILPs*, FOXO target genes and *foxo*. (A-C) Whole larvae mRNA levels measured by qRT-PCR in control (*hml > +*) versus TFAM RNAi (*hml > UAS-TFAM-RNAi*) larvae at 96 hrs AEL. Data are represented as mean ± SEM, with individual data points plotted as symbols (*p < 0.05 and ns, not significant, unpaired t-test). n (# of samples) = 6 (control) and 6 (TFAM RNAi).

**Figure S3:**
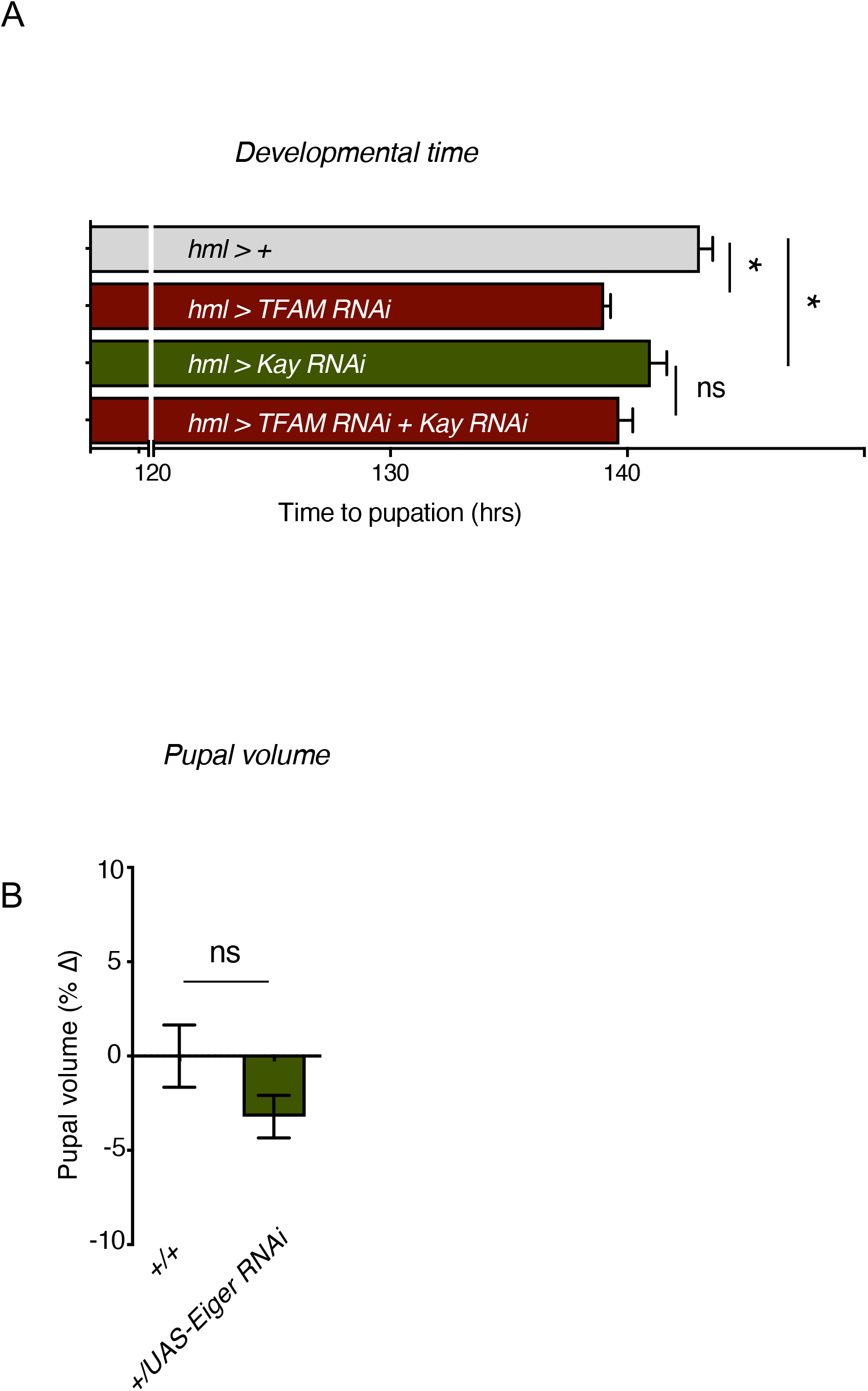
Hemocyte specific knock down if JNK pathway components suppresses systemic growth (related to Fig. 5 and 6) (**A**) Time to pupation was measured in control (*hml > +*), TFAM RNAi (*hml > UAS-TFAM RNAi*), Kay RNAi (*hml > UAS-Kay RNAi*) and TFAM RNAi + Kay RNAi (*hml > UAS-TFAM RNAi + UAS-Kay RNAi*) larvae. Data are represented as mean ± SEM, with individual data points plotted as symbols (*p < 0.05 and ns, not significant, unpaired t-test). n (# of samples) = 278 (control), 460 (*TFAM RNAi*), 128 (*Kay RNAi*), and 126 (*TFAM RNAi + Kay RNAi*). (**B**) Relative change in pupal volume was calculated based on the average value of control (+/+) animals. Data are presented as mean +/− SEM (*p < 0.05, Mann-Whitney U test) for controls and Eiger RNAi lines (*+/UAS-Eiger-RNAi*). n (# of pupae) = 76 (control) vs 148 (Eiger RNAi).

